# Uncovering Hidden Enhancers Through Unbiased *In Vivo* Testing

**DOI:** 10.1101/2022.05.29.493901

**Authors:** Brandon J. Mannion, Marco Osterwalder, Stella Tran, Ingrid Plajzer-Frick, Catherine S. Novak, Veena Afzal, Jennifer A. Akiyama, Sarah Barton, Erik Beckman, Tyler H. Garvin, Patrick Godfrey, Janeth Godoy, Riana D. Hunter, Momoe Kato, Michal Kosicki, Anne N. Kronshage, Elizabeth A. Lee, Eman M. Meky, Quan T. Pham, Kianna von Maydell, Yiwen Zhu, Javier Lopez-Rios, Diane E. Dickel, Axel Visel, Len A. Pennacchio

## Abstract

Transcriptional enhancers are a predominant class of noncoding regulatory elements that activate cell type-specific gene expression. Tissue-specific enhancer-associated chromatin signatures have proven useful to identify candidate enhancer elements at a genome-wide scale, but their sensitivity for the comprehensive detection of all enhancers active in a given tissue *in vivo* remains unclear. Here we show that a substantial proportion of *in vivo* enhancers are hidden from discovery by conventional chromatin profiling methods. In an initial comparison of over 1,200 *in vivo* validated tissue-specific enhancers with tissue-matched mouse developmental epigenome data, 14% (n=286) of active enhancers did not show canonical enhancer-associated chromatin signatures in the tissue in which they are active. To assess the prevalence of enhancers not detectable by conventional chromatin profiling approaches in more detail, we used a high throughput transgenic enhancer reporter assay to systematically screen over 1.3 Mb of mouse genomic sequence at two critical developmental loci, assessing a total of 281 consecutive 5kb regions for *in vivo* enhancer activity in mouse embryos. We observed reproducible enhancer-reporter activity in 88 tissue-specific elements, 26% of which did not show canonical enhancer-associated chromatin signatures in the corresponding tissues. Overall, we find these hidden enhancers are indistinguishable from marked enhancers based on levels of evolutionary conservation, enrichment of transcription factor families, and genomic positioning relative to putative target genes. In combination, our retrospective and prospective studies assessed only 0.1% of the mouse genome and identified 309 tissue-specific enhancers that are hidden from current chromatin-based enhancer identification approaches. Our findings suggest the existence of tens of thousands of active enhancers throughout the genome that remain undetected by current chromatin profiling approaches and are an unappreciated source of additional genome function of import in interpreting growing whole human genome sequencing data.

## Introduction

The importance of distant-acting enhancers in the temporal and spatial control of human gene expression is well established^1–4^. Proper transcriptional regulation by enhancers, which are particularly enriched near developmentally important genes, enables normal organismal development and function^5–7^. The initial characterization of enhancers was enabled by pioneering molecular studies of individual loci such as locus control regions at the β-globin locus^8–10^, the availability of initial noncoding comparative genomic information from species such mouse, rat, and pufferfish^11–13^, and powerful genomic approaches including ChIP-chip and subsequent next generation sequencing techniques^14–16^.

Dedicated genomic efforts such as ENCODE have sought to systematically identify enhancers via suitable *in vitro* and *in vivo* approaches^17^. Remarkably, while the human genome contains only ∼20,000 protein encoding genes, these studies identified on the order of one million putative enhancers^18^. For example, a ChIP-Seq study that examined the enhancer-associated mark H3K27ac on a panel of 12 tissues isolated through daily sampling during mouse development uncovered ∼200,000 candidate enhancers^19^. Chromatin accessibility (by ATAC-seq), H3K4me1, and H3K27ac are utilized as canonical enhancer-associated chromatin signatures^16,20–22^. However, the accuracy and practical utility of these data sets critically depends on the correlation of the marks examined with true *in vivo* activity, which can be assessed in transgenic reporter assays^23^. For instance, enhancer validation efforts in mouse *in vivo* assays revealed the potential for substantial false positives in these putative H3K27ac derived enhancer datasets^19^. In contrast, the comprehensiveness (false negatives rates) of these enhancer catalogs remains unknown, which we examine as a central goal in the present study^24^.

To assess the prevalence and characteristics of enhancers potentially missed in current datasets, we use pre-existing VISTA validated enhancers and the ENCODE chromatin catalog to retrospectively compare reproducible VISTA tissue-specific activity with their chromatin-based signatures from identical tissues *in vivo*^*19,25*^. We then performed large-scale transgenic enhancer assays^26^ for the unbiased tiling of over 1.3 Mb of the mouse genome in an attempt to replicate our initial observations and uncover additional hidden enhancers within mammalian genomes. Indeed, through these retrospective and prospective studies, we show that hidden enhancers exist within our genome and will require a combination of improved genome-wide profiling methods and *in vivo* experimentation to identify and characterize them.

## Results

### Many *in vivo* enhancers show no canonical enhancer marks

As an initial exploration of the comprehensiveness of chromatin-based enhancer mapping strategies, we used the VISTA Enhancer Browser database (https://enhancer.lbl.gov)^25^ to retrospectively assess the relationship between enhancer-associated chromatin marks and validated enhancer activity *in vivo*. To date, this resource includes over 3,200 human and mouse elements that have been tested for enhancer-reporter activity, primarily at mouse embryonic day 11.5 (E11.5), a stage when multiple developing tissues (*e*.*g*., limb, craniofacial structures) can be assessed through whole-mount imaging in mice and compared with their functional counterparts in humans. We focused on 1,272 validated enhancers driving expression in one or more anatomical structures including forebrain (n=450); midbrain (398); hindbrain (366); craniofacial region (262); limb (304); and heart (272). We compared these data to perfectly matched epigenetic tissue data that includes ATAC-seq, H3K27ac ChIP-seq, and H3K4me1 ChIP-seq (**Table S1**). For each of the six tissues, we examined the presence of canonical enhancer-associated chromatin signatures at each positive element’s endogenous site (**Fig. 1a-b, Table S2**).

**Figure 1.**
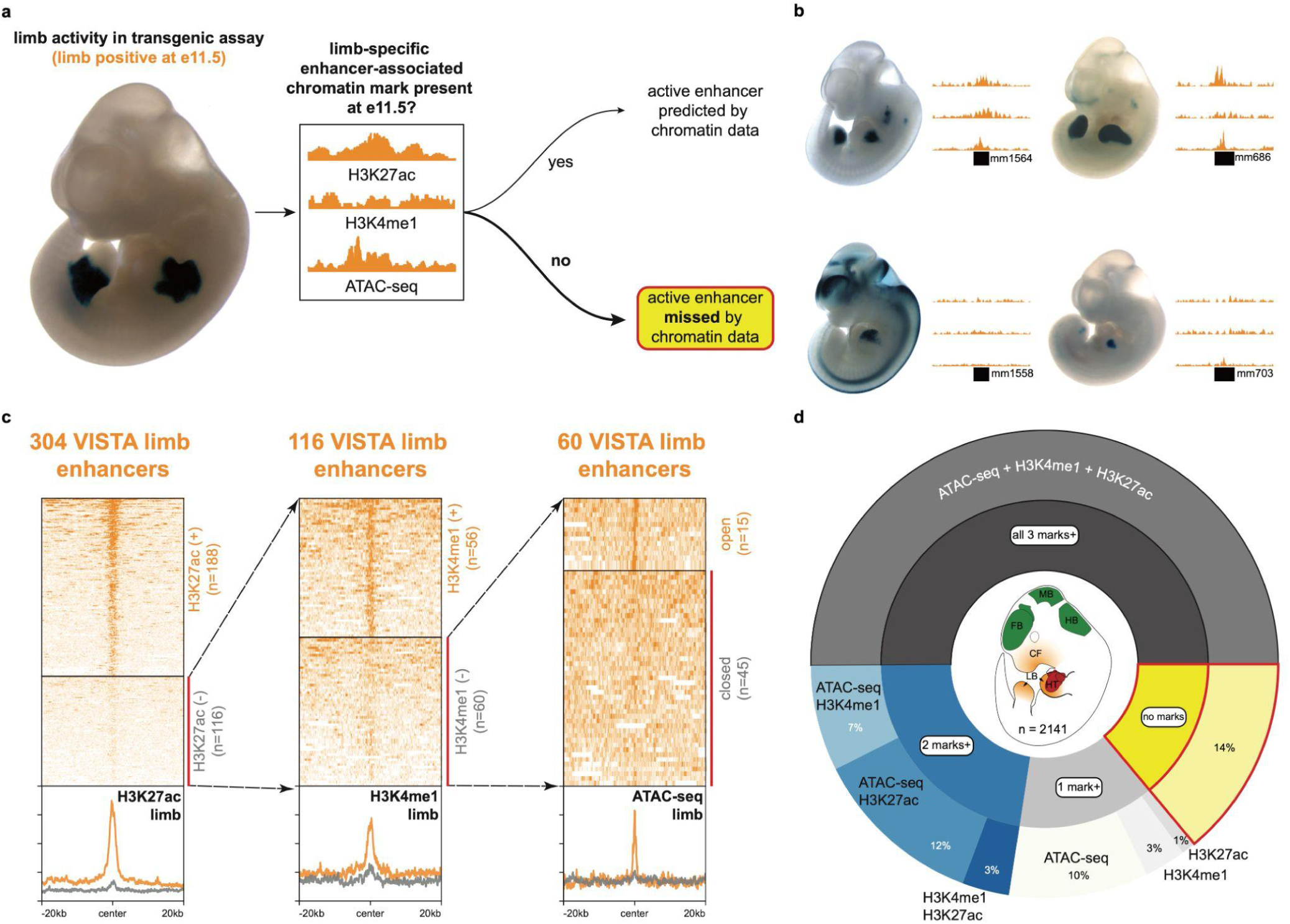
Mouse *in vivo* enhancers without canonical enhancer-associated chromatin marks. **(a)** Approach to retrospectively identify active enhancers without tissue-specific enhancer-associated chromatin marks. **(b)** Examples of active limb enhancers with (top row) and without (bottom row) enhancer-associated chromatin marks in dissected tissue. See **Fig. S1** for examples of active enhancers without these marks in other tissues. **(c)** Chromatin profiles of active limb enhancers with and without H3K27ac (ChIP-seq), H3K4me1 (ChIP-seq), or open chromatin (ATAC-seq). See **Fig. S2** for another example of chromatin mark filtering for forebrain enhancers. **(d)** Proportion of VISTA enhancers across six tissues (forebrain, midbrain, hindbrain, craniofacial structure, limb, heart) with and without enhancer-associated chromatin marks. We focused on the VISTA enhancers with activity (“positive” elements) in the above six tissues, however we also observed enhancers that are active in other tissues at E11.5 (**Table S3**). Active enhancers without any of these chromatin marks are in yellow.

For example, for the 304 VISTA limb enhancers, we found 116 (38%) do not have a limb-specific H3K27ac enhancer-associated mark (**Fig. 1c**). In addition, of these 116 limb enhancers lacking H3K27ac marks, 60 (20%) also lack an H3K4me1 mark. Finally, 45 of these limb enhancers (15% of VISTA limb-positive elements) are completely lacking any of the three enhancer-associated chromatin marks (limb-specific H3K27ac, ATAC-seq, or H3K4me1). Across the six tissues examined, these “hidden” enhancers represent 9% to 25% of VISTA enhancers (**Fig. S3**). Overall, we found that 50% (1028) of tissue-specific VISTA enhancers have all three marks, 22% (461) have at least two marks, 13% (277) have only one of the three marks, and 14% (286) are hidden enhancers without any of the three marks in the corresponding tissue. The relative proportions of enhancer mark categories are similar across the six considered tissues. We observed “hidden” enhancers in all developing tissues (*e*.*g*., forebrain, craniofacial structure, heart) that we assessed at E11.5, suggesting their existence is a general phenomenon.

### Mouse *in vivo* tiling assay uncovers additional hidden enhancers

Since many of the enhancers reported in the literature and VISTA database were found through chromatin signature-guided enhancer discovery screens, retrospective intersections are likely to underestimate the proportion of enhancers lacking canonical chromatin signatures. To assess this phenomenon in a more unbiased manner, we selected two separate loci (*Gli3*; *Smad3/Smad6*) to test the enhancer activity of over 281 overlapping elements regardless of their chromatin state. The *Gli3* gene encodes a transcription factor involved in multiple pathways that are involved in the development of the limb, face, and nervous system^27–29^. Apart from *Gli3* itself, the flanking region considered for tiling is generally depleted of other genes, which includes a gene desert upstream of *Gli3* that spans over 800kb^30^. Dozens of regions (n=38) across the locus are predicted to be enhancers based on tissue-specific H3K27ac (**Table S4, Fig. 2a**) and prior limited candidate enhancer studies within this locus demonstrated limb and brain-related enhancer activity in E11.5 mouse^31,32^. Additionally, we assessed a separate locus for unbiased tiling that encompasses the *Smad3* and *Smad6* genes (**Fig. S4**). As with the *Gli3* locus, the *Smad3*/*Smad6* locus considered for tiling also includes several (n=86) H3K27ac-marked regions (**Table S4**). While *Smad3* is broadly expressed in all six of the tissues examined in this study at mouse E11.5, expression of *Smad6* is highest in mouse E11.5 heart (**Fig. S5**), which is supported by earlier characterizations of *Smad6* in cardiovascular development^33^. For both loci, we used these earlier findings and the aforementioned H3K27ac-marked regions as positive benchmarks to compare with the tiling elements, which we designed as overlapping elements (average size ∼5kb) with boundaries chosen to fully capture complete H3K27ac-enriched regions where possible and tested the sequences in a mouse *in vivo* transgenic assay (**Fig. 2b-c**). Altogether, these 281 tested elements span over 1.3 Mb of the mouse genome.

**Figure 2.**
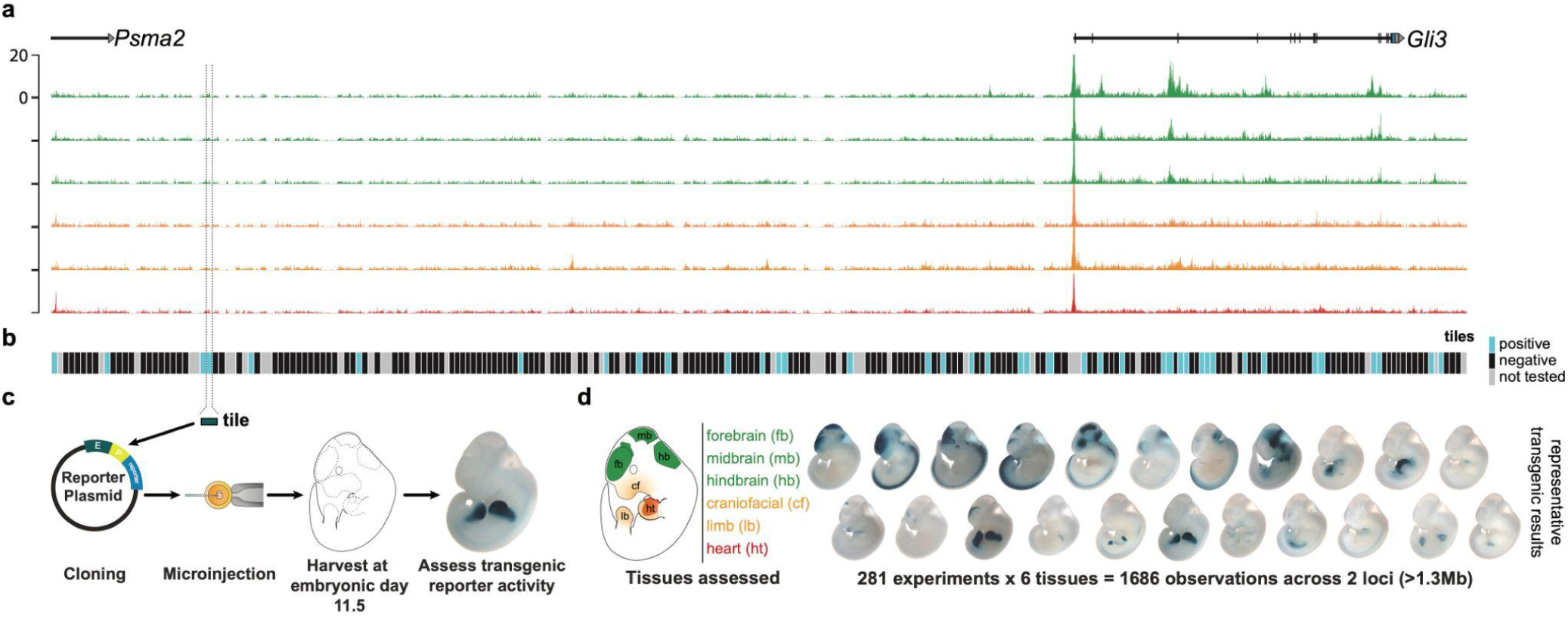
Systematic tiling for the unbiased identification of mouse in vivo enhancers. **(a)** *Gli3* locus with H3K27ac ChIP-seq data (ENCODE) for six tissues. **(b)** Elements (∼5kb in size and overlapping with adjacent elements) designed for the unbiased tiling assay. Elements that were tested and that had reproducible enhancer-reporter activity in the mouse *in vivo* transgenic assay are shaded blue. We observed 63 active enhancers at mouse embryonic day 11.5 (E11.5) with tissue-specific activity. Of these 63 enhancers, 36 show reproducible enhancer-reporter activity in multiple tissues at E11.5. Elements without reproducible activity are shaded black. Elements not successfully tested are shaded gray. **(c)** Approach for testing each tile in the mouse *in vivo* transgenic assay. (**d**) Depiction of tissues that were checked for reproducible enhancer-reporter activity and example transgenic results from tiling across the *Gli3* and *Smad3/Smad6* loci (see **Fig. S4** for the *Smad3/Smad6* locus).

We observed that 63 of 281 tested elements showed reproducible enhancer activity at mouse embryonic day 11.5 (E11.5) in at least one tissue (**Fig. 2b, Fig. S4**). Similar to the retrospective VISTA study, we focused on six tissues (forebrain, midbrain, hindbrain, craniofacial structure, limb, heart) to compare *in vivo* activities with tissue-matched chromatin data. A majority (43) of active enhancers from the tiling assay show LacZ reporter activity in E11.5 tissues that recapitulate the known expression patterns of the nearest genes within the tiled loci (**Fig. S6**).

We next assessed the relationship between experimental enhancer data in transgenic reporter assays and epigenomic data (H3K27ac, ATAC-seq, H3K4me1) from the six tissues of interest (**Table S2**). Of the 63 active enhancers discovered in functional mouse tiling assays, which represent 88 tissue-specific enhancer activities, we observed that 23 (26%) were hidden (*i*.*e*., they lack enhancer-associated chromatin marks in their active tissue(s)). From the unbiased tiling, we observed that hidden enhancers represent a larger proportion of active enhancers (26%) relative to the retrospective VISTA enhancer study (14%) described above (**Fig. 3a**). We identified hidden enhancers in all 6 tissues under investigation, such as forebrain (n=2, least) and hindbrain (n=7, most) (**Fig. 3b**). In addition to exploring false-negatives in enhancer identification, these data also allowed us to explore false-positives. Of those 141 regions that were both marked with tissue-specific H3K27ac and tested in the transgenic assay, 23% did validate as active enhancers in their corresponding tissue(s). For either locus, high-ranking tissue-specific H3K27ac peaks, *i*.*e*., those regions with a higher enrichment of mapped reads vs. background, more often validated as active enhancers relative to middle- or low-ranking peaks which is consistent with previous reports^18,19^ (**Fig. S6, Table S5**). Altogether, our retrospective VISTA study and unbiased systematic experimental testing uncovered 309 tissue-specific hidden enhancers, supporting the existence of substantial numbers of missing enhancers genome-wide.

**Figure 3.**
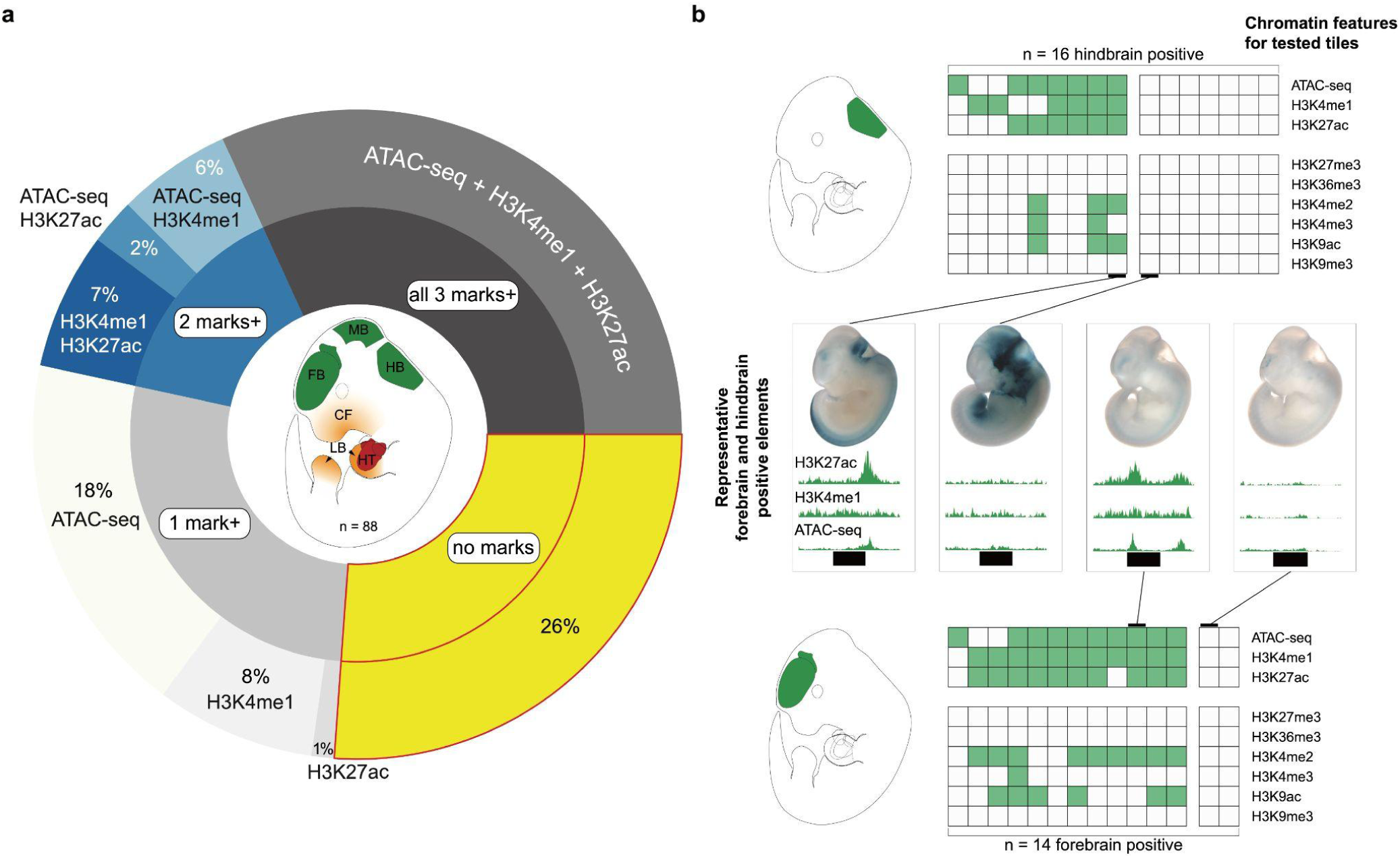
Active enhancers from unbiased tiling with and without enhancer-associated chromatin marks. **(a)** Proportion of active enhancers from unbiased tiling with and without enhancer-associated chromatin marks. Active enhancers without any of these chromatin marks are in yellow. **(b)** Example of active hindbrain (n=16) and active forebrain enhancers (n=14) from the tiling assay. Columns within the square table represent a tested element (a region with genomic coordinates), whereas rows within a single column represent the chromatin feature (shaded green if a peak in the given chromatin feature is present) for that particular element. The square tables are split into two main categories, those elements with at least one of the considered enhancer-associated chromatin marks present (left) and those without any of the three considered enhancer-associated chromatin marks (right). Additional chromatin data depicted show that a portion of hidden enhancers are absent of all chromatin marks assayed by ENCODE (**Fig. S7**). Representative transgenic results (2 for hindbrain; 2 for forebrain) are depicted as well as the chromatin profile for the relevant element.

### Hidden enhancers are indistinguishable from their marked counterparts

We next assessed the properties of hidden versus marked enhancers in an attempt to explain their functional differences. In particular, within each tiling locus we did not find specific transcription factor families that were enriched in hidden enhancers relative to marked enhancers (**Table S6**). Similarly, by functional enrichment analysis there are no significant biological processes or phenotypes that distinguish hidden enhancers from their marked counterparts (**Table S7**). From the VISTA retrospective study and the unbiased tiling, both hidden and marked enhancers have similar levels of evolutionary conservation (**Fig. S8**). Further, we checked the positional distribution of either enhancer group within their topological associated domains (TADs) and, based on mESC TADs^34^, we did not find hidden enhancers to be enriched in specific positions relative to their marked counterparts (**Fig. S9**). Based on these findings, we did not identify any robust and specific properties of hidden enhancers that could potentially enable their genome-wide identification.

### Hidden enhancers cannot be identified by existing epigenomic data

Given the absence of canonical enhancer marks in embryonic mouse tissue-derived epigenomic data sets, we examined if complementary epigenomic data types offer potential avenues for the discovery of these hidden enhancers. We first evaluated if hidden enhancer activity at E11.5 could be the outcome of residual LacZ reporter activity resulting from expression that occurred at an earlier developmental stage. We found that of the 309 tissue-specific hidden enhancers assayed at E11.5, 173 (56%) have enhancer-associated chromatin marks at an earlier stage, *i*.*e*., H3K27ac and/or H3K4me1 at embryonic day 10.5 (E10.5).

Next, we checked if available single-cell chromatin data could resolve enhancer-associated chromatin marks around hidden enhancers that might have been missed from standard chromatin data derived from bulk tissue preparations. Of the six tissues for which we compared transgenic enhancer-reporter activity with the corresponding mouse tissue chromatin data from ENCODE, two tissues (forebrain, hindbrain) have single nucleus ATAC-seq (snATAC-seq) data across early mouse development^35,36^. For the 50 hidden enhancers that are either forebrain or hindbrain enhancers, only 8 (16%) could be identified via available corresponding single cell data.

Finally, since 236 (83%) of the 286 hidden enhancers identified in the VISTA retrospective study are human sequences tested in mouse transgenic enhancer assays, we checked if available human tissue-matched epigenomic data would have predicted any of these hidden enhancers. For 112 human-derived hidden enhancers that did not have enhancer-associated chromatin marks either in earlier (E10.5) or in single-cell chromatin data, 49 were assessable with available tissue-matched, similar-staged human chromatin data from craniofacial, heart, and limb bud tissues. Only 10 (20%) showed enhancer marks in available human chromatin data. Altogether through these analyses, 118 (38%) of the originally identified hidden enhancers could not be identified despite at least two of the three complementary data types being available (**Fig. 4**), further supporting the wide-spread genome wide occurrence of hidden enhancers.

**Figure 4.**
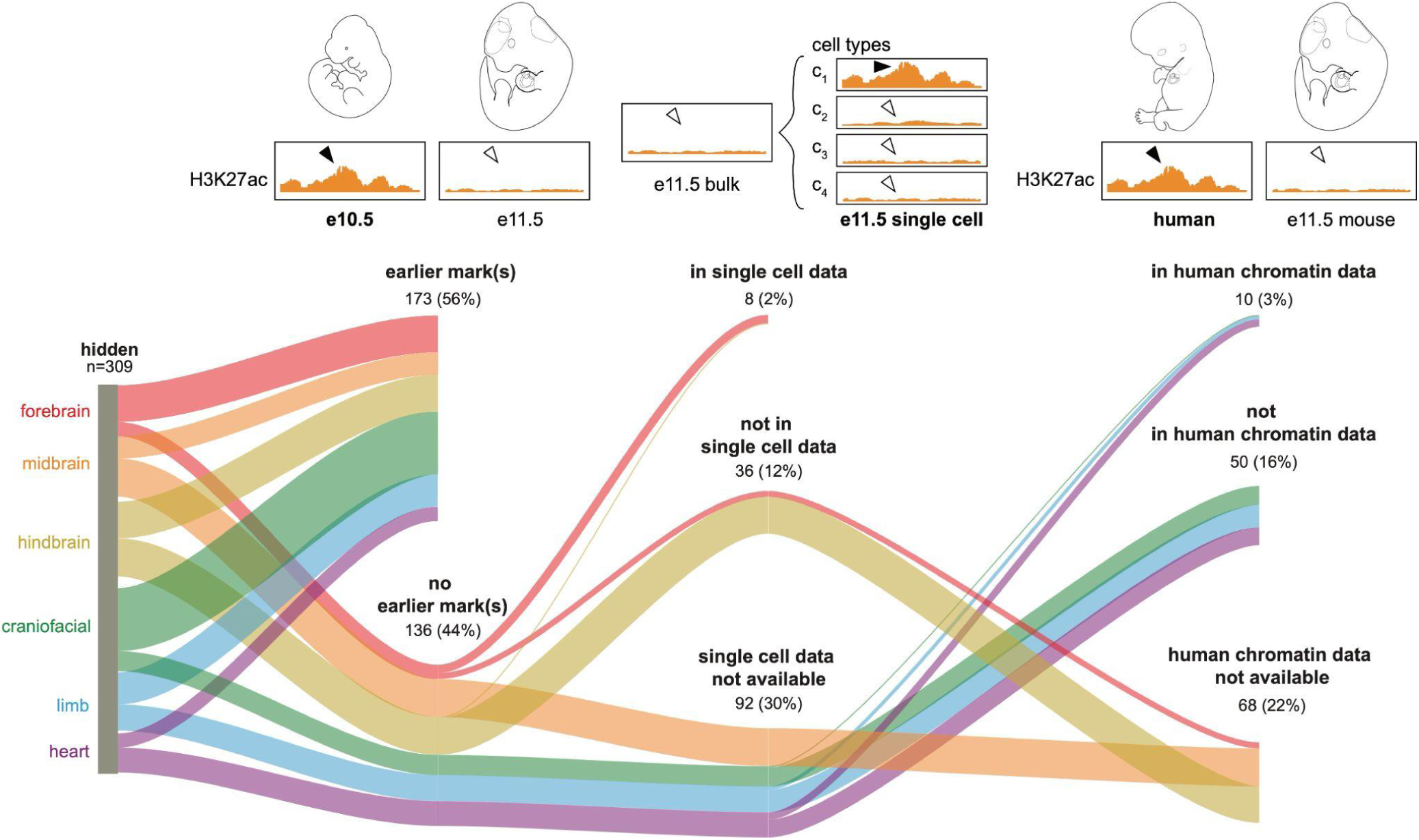
Hidden enhancers cannot be fully recovered from alternative chromatin data. **(a)** Hidden enhancers that are active at E11.5 either do or do not have earlier enhancer-associated chromatin marks at E10.5. **(b)** Hidden enhancers based on bulk tissue chromatin data either are or are not annotated with enhancer-associated chromatin marks from available single cell chromatin accessibility data. **(c)** Hidden enhancers tested in the mouse *in vivo* transgenic assay either do or do not have corresponding enhancer-associated chromatin marks in available human chromatin data.

## Discussion

In this study, we report the existence of hundreds of hidden enhancers in the human/mouse genome that lack well-established enhancer-associated marks from their precise tissue of activity. This includes a retrospective analysis of over 1,200 *in vivo* validated tissue-specific enhancers in VISTA as well as an unbiased prospective study of ∼280 elements representing consecutive tiles across 1.3 Mb of the mouse genome. We found a moderate portion of these hidden enhancers can be identified by considering complementary data either from other time points, single cell chromatin measurements, or other species. While public single cell chromatin accessibility data are currently limited to a few tissues in mice^35,36^, their generation from different additional tissues (*e*.*g*., face, limb, heart) should provide higher resolution for rare cell populations than bulk data in which tissue-specific enhancer marks might be missed. Additionally, human chromatin data within a similar developmental window are currently available only from a few tissues (*i*.*e*., heart^37^, face^38^, limb bud^39^), and future characterization of other similarly staged human tissues should facilitate cross species comparisons of enhancer-associated chromatin marks and *in vivo* enhancer activity. Altogether, we could not identify sequence, genomic, or other epigenomic properties that could distinguish hidden enhancers from their marked counterparts. However, alternative features (across differing experimental systems) that include chromatin architecture and enhancer-promoter communication^40–42^, transcriptional output^43^, and non-canonical histone marks^44^ may provide additional insights on hidden enhancer activity. These findings support the need for novel experimental and/or interpretable computational approaches that harness and combine available data for more systematic genome-wide enhancer identification.

In addition to uncovering hidden enhancers, which represent false-negatives in existing epigenomic data, this study also allowed us to further investigate false-positives in these datasets (*i*.*e*., putative enhancers identified in epigenomic data but not validated in transgenic assays). Our finding that high-ranking H3K27ac peaks across the tiling loci had higher *in vivo* validation rates than lower-ranking peaks is consistent with previous candidate enhancer studies based on similarly ranked chromatin data^19^. Our study further confirms both the importance of being cautious for the interpretation of possible missing information (hidden enhancers) and the consideration of false-positives in otherwise powerful epigenomic enhancer catalogs.

This prospective tiling study implemented a recently scaled transgenic assay^26^ to systematically test elements for mouse *in vivo* enhancer activity across over 1.3 Mb of a mammalian genome. With the current setup of the transgenic assay, we evaluated only the sufficiency of candidate elements for enhancer-reporter activity at a non-endogenous site. Nevertheless, the enhancer elements (over 280 ∼5kb tiles) that we have characterized provide validated targets to explore further the sequence and/or epigenetic properties that are important for enhancer function. Across the two tiling loci we found over 80 tissue-specific enhancers, around 26% of which are hidden enhancers. We focused on only six tissues from which we could compare between tissue-matched chromatin properties and mouse *in vivo* data at E11.5, yet there are vast numbers of other tissues and developmental time points relevant for enhancer identification^45^. With some estimates of hundreds of thousands to nearly one million candidate enhancers in mammalian genomes, one might speculate from our tiling study that there are tens of thousands of additional enhancers unaccounted for by current approaches^18^. As sequencing expands to cover the full range of human tissues, diversity, environmental perturbations, and as related technologies provide even higher resolution approaches to probe gene regulatory activity, we can expect to better understand and annotate the unique characteristics of hidden enhancers and their functional significance in transcriptional regulation.

## Materials and Methods

### ENCODE mouse chromatin and RNA-seq data

Processed mouse chromatin data^19^ (ATAC-seq; ChIP-seq for H3K27ac, H3K4me1, H3K4me2, H3K4me3, H3K27me3, H3K9ac, H3K36me3, H3K9me3) and RNA-seq data^46^ were downloaded from the ENCODE resource portal (https://www.encodeproject.org/). Details on the generation and processing of these data are available here: https://www.encodeproject.org/pipelines/. See Supplementary Table 1 for a listing of all the bulk tissue mouse data used for chromatin intersections or tissue expression analyses.

### VISTA enhancers

Human and mouse candidate enhancers were tested in a mouse *in vivo* transgenic reporter assay, as previously described^26^ (see also “Locus selection for tiling and mouse in vivo enhancer validation”). Candidate enhancers were assessed for reproducible enhancer-reporter activity in forebrain, midbrain, hindbrain, craniofacial structures (*e*.*g*., branchial arches; nose; facial mesenchyme), limb, and heart. The genomic coordinates (assembly mm10) of these elements were downloaded from the VISTA Enhancer Browser (https://enhancer.lbl.gov/)^25^. Human elements were lifted over from hg38 to mm10 via the UCSC liftOver tool using *minMatch=0*.*1*^*47*^.

### Chromatin intersections and hidden enhancer identification

Mouse *in vivo* validated elements from both the VISTA Enhancer Browser and the tiling assay were intersected with tissue-specific ENCODE chromatin data via bedtools^48^ (v2.29.0) to check for the presence or absence of enhancer-associated chromatin signatures (*e*.*g*., tissue-specific mouse E11.5 peaks from ATAC-seq, H3K27ac ChIP-seq, and/or H3K4me1 ChIP-seq data) within each elements’ genomic coordinates. Elements with reproducible enhancer-reporter activity (positive elements) but without any of the three enhancer-associated chromatin signatures in the relevant tissue(s) were designated as hidden enhancers. Positive elements with any (up to all) of the three enhancer-associated chromatin signatures were considered marked enhancers. Positive elements were also checked for overlap with other chromatin features available from mouse ENCODE: DNase-seq, H3K27me3 ChIP-seq, H3K36me3 ChIP-seq, H3K4me2 ChIP-seq, H3K4me3 ChIP-seq, H3K9ac ChIP-seq, and H3K9me3 ChIP-seq. Both mouse embryonic days 10.5 (E10.5) and 11.5 (E11.5) data were used for the above analyses.

### Locus selection for tiling and mouse *in vivo* enhancer validation

Coordinates used for the *Gli3* locus are chr13:14,626,494-15,785,614 (mm10). Coordinates used for the *Smad3/Smad6* locus are chr9:63,685,831-64,099,907 (mm10). Tiling elements approximately 5kb in size (with overlap to adjacent tiling elements) were cloned into the pCR4-Shh::lacZ-H11 vector (Addgene plasmid #139098), which includes the mouse Shh promoter, the *LacZ* gene for enzymatic, colorimetric readout, and flanking homology arms that enable site-specific integration at the H11 locus^26^. A mixture of the reporter construct, Cas9 protein, and sgRNAs were transferred by microinjection into the pronucleus of mouse embryos (FVB strain) and then transferred to the uterus of pseudopregnant females (CD-1 strain). Transgenic embryos were then collected at mouse embryonic day 11.5 (E11.5) for LacZ staining and the assessment of enhancer-reporter activity in several developing tissues (*e*.*g*., forebrain, midbrain, hindbrain, craniofacial, heart, and limb). For more detailed steps and information on the workflow that spans cloning, mouse colony management, microinjection, and embryo staining, refer to the recently published protocol^49^. Transgenic embryos for each tiling element were assessed for reproducible enhancer-reporter activity in separate, independent embryos. Genomic coordinates, transgenic embryo images, and tissue annotations for each element are available on the VISTA Enhancer Browser (https://enhancer.lbl.gov).

### Enhancer mark *in vivo* validation rates

For each tissue, we selected H3K27ac peaks that were at least 1,000bp from an annotated transcription start site (GENCODE)^50^ and that also did not overlap annotated exons, UTRs, and stop codons. H3K27ac peaks within 2,500bp of each other were merged and the maximum -log_10_(q-value) among these peaks was kept. To assess the mouse *in vivo* validation rates of the H3K27ac enhancer-associated chromatin mark, the final peak list was ranked by highest -log_10_(q-value) and then compared with the mouse *in vivo* transgenic result of the overlapping element (genomic region).

### Evolutionary conservation

PhastCons scores were downloaded from the UCSC Genome Browser at https://hgdownload.cse.ucsc.edu/goldenPath/mm10/phastCons60way/. phastCons scores were calculated for each element (mean across region) and used to compare the levels of evolutionary conservation between different categories of tested elements (*e*.*g*., hidden enhancers vs. marked enhancers). The Kolmogorov-Smirnov test was used to assess potential differences in phastCons distributions between the considered enhancer categories.

### Distribution of elements within topologically associated domains

Topologically associated domains (TADs) called from mouse embryonic stem cell Hi-C^34^ were downloaded (http://chromosome.sdsc.edu/mouse/hi-c/download.html) and domain coordinates were lifted over from assembly mm9 to mm10 via the UCSC liftOver tool using minMatch=0.95. Human elements (tested in the mouse *in vivo* system) were lifted over from hg38 to mm10 via the UCSC liftOver tool using *minMatch=0*.*1*. The distance (bp) of each enhancer (hidden and marked) from the nearest TAD boundary was calculated and then normalized to the size of the given TAD (% TAD length). The distributions of these normalized positions were plotted (per tissue and across all tissues), and the Kolmogorov-Smirnov test was used to compare for potential differences between hidden and marked enhancers.

### Additional epigenomic data

Publicly available single cell chromatin accessibility data from mouse E11.5 forebrain (GSE100033)^35^ and mouse E11.5 cerebellum (https://apps.kaessmannlab.org/mouse_cereb_atac/)^36^ were used to compare differences between bulk tissue and single cell assays in the resolution of enhancer-associated chromatin signatures, *i*.*e*., if there were open chromatin regions absent in bulk chromatin data but detected in single cell data. Human chromatin data from approximately stage-matched limb bud (GSE42413)^39^, heart (GSE137731)^37^, and face (GSE97752)^38^ were used to evaluate if hidden enhancers from human sequence could be identified with these complementary data.

### Functional enrichment analysis

GREAT^51^ version 4.0.4 (http://great.stanford.edu/public/html/) was used to assess the enrichment of biological ontologies in hidden enhancers, via the basal plus extension setting (5,000bp upstream, 1,000bp downstream, distal up to 1Mbp).

### Transcription factor motif analysis

HOMER^52^ version 4.10 was used to assess enrichment of both known and *de novo* motifs within hidden enhancers, via *findMotifsGenome*.*pl* and the following parameters: -size given -len 8,9,10,12,14 -bg <background file = all positive VISTA enhancers>.

## Supplementary Figures

**Figure S1.**
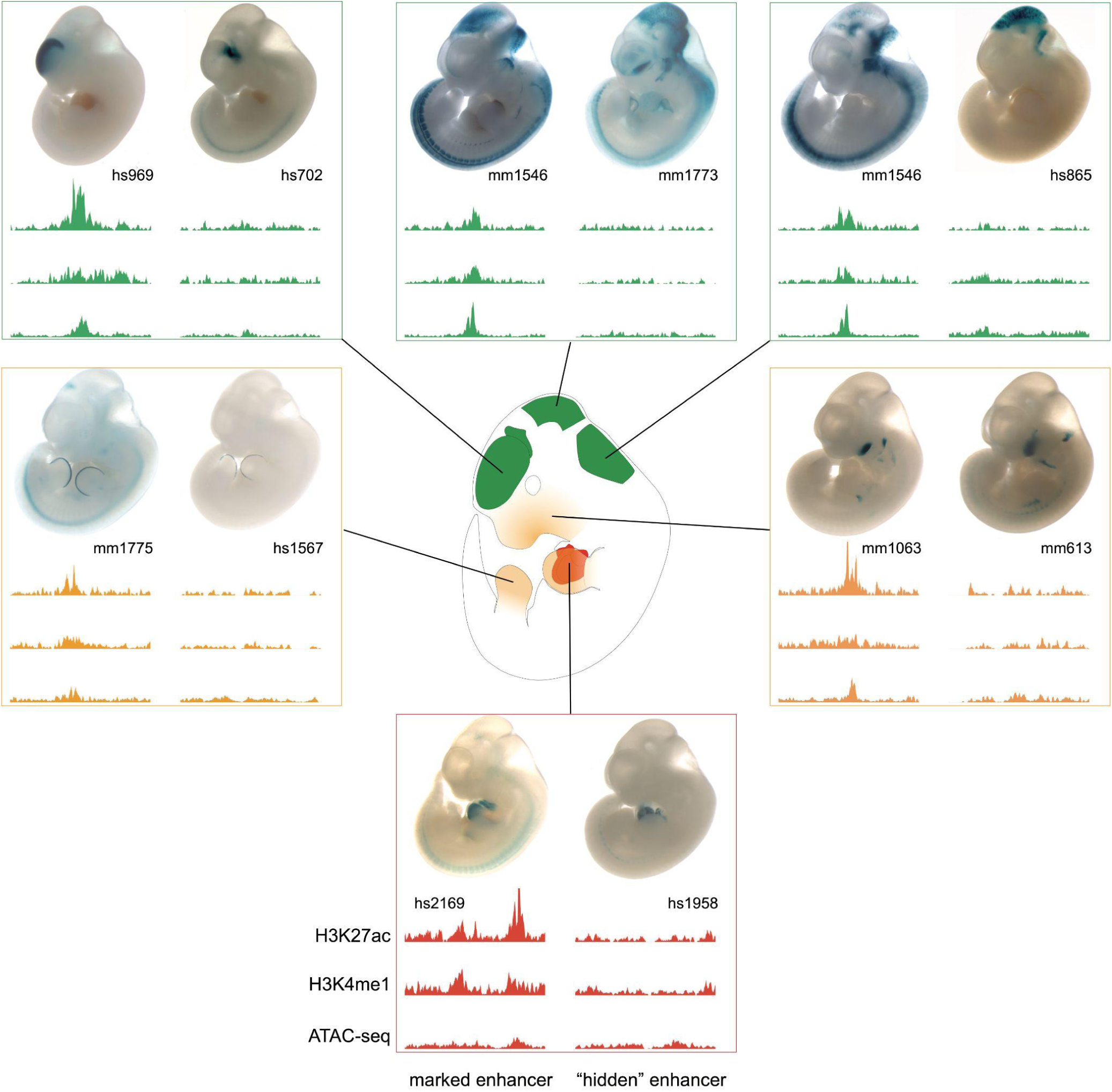
Mouse *in vivo* enhancers with and without canonical enhancer-associated chromatin marks. Representative transgenic result (mouse E11.5 embryos) displayed above tissue-specific chromatin profile for each tested element (VISTA ID provided). For each of the 6 considered tissues, an active enhancer with canonical enhancer-associated chromatin marks (left) is displayed alongside an active enhancer without canonical enhancer-associated chromatin marks (right). Mouse tissue- and stage-matched H3K27ac ChIP-seq, H3K4me1 ChIP-seq, ATAC-seq are from ENCODE^19^.

**Figure S2.**
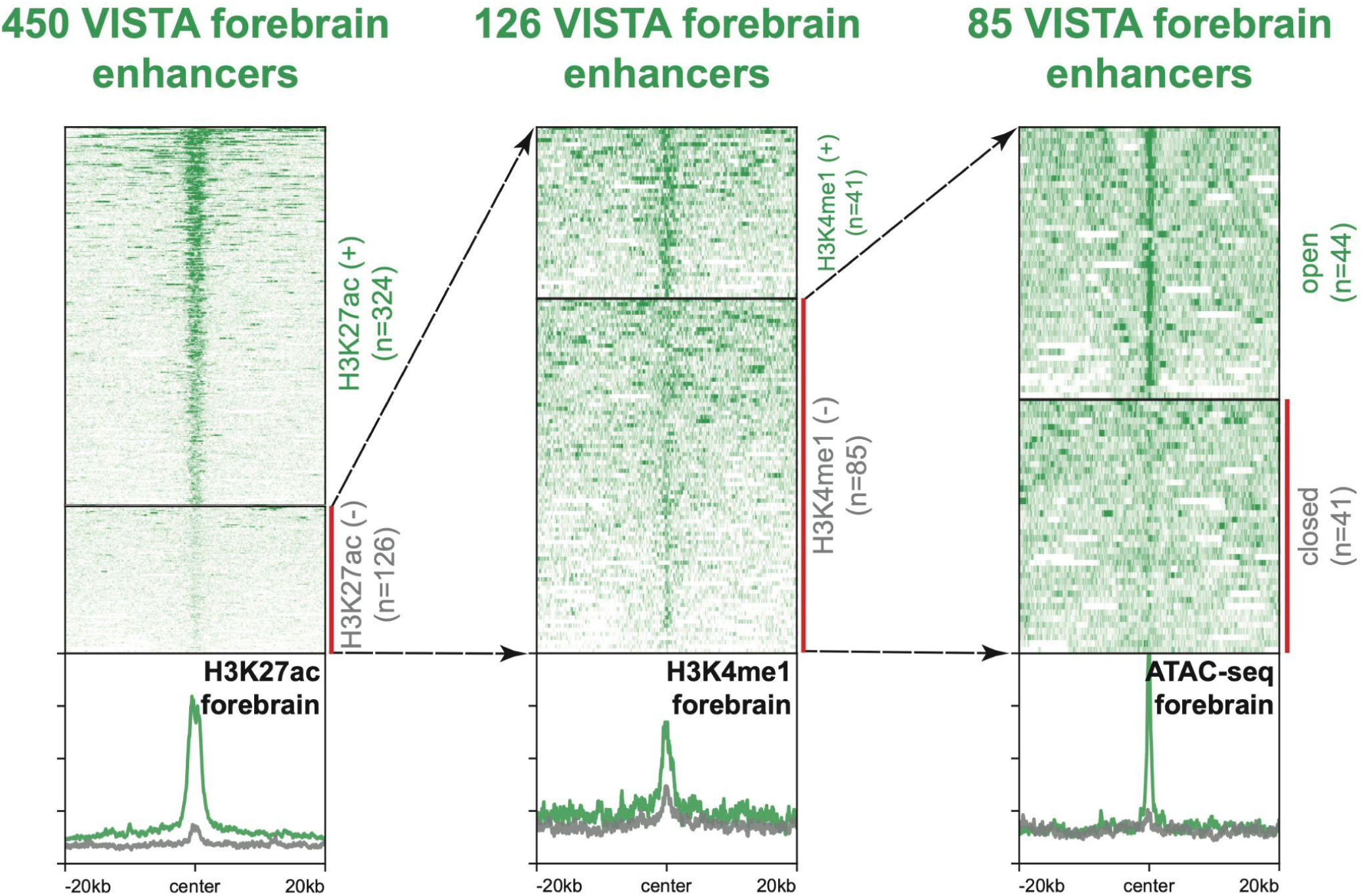
Chromatin profiles of active forebrain enhancers with and without H3K27ac, H3K4me1, and ATAC-seq (open chromatin). Forebrain enhancers from the VISTA Enhancer browser stratified across three canonical enhancer-associated chromatin marks. Processed mouse chromatin data are from ENCODE^19^.

**Figure S3.**
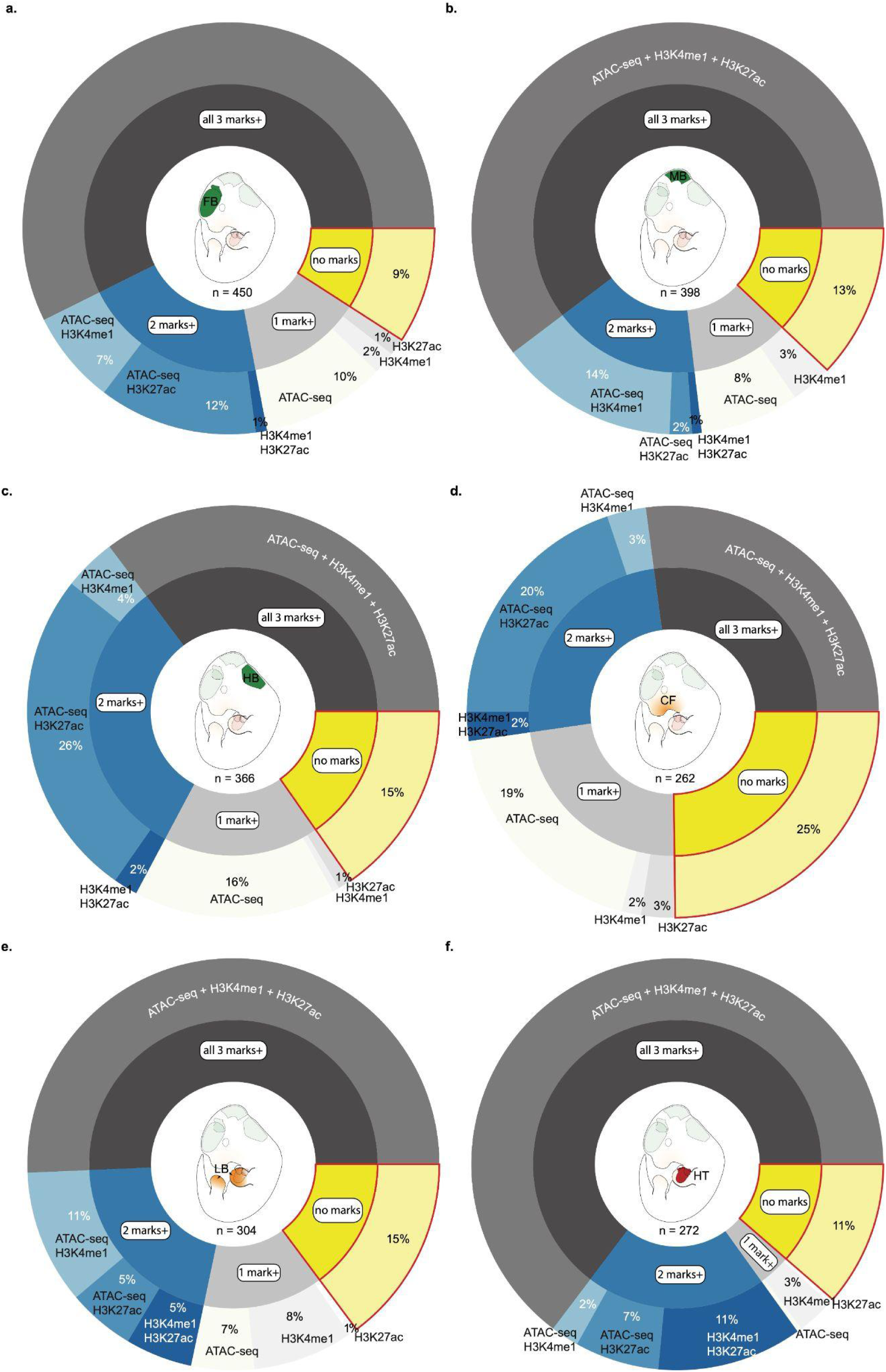
Proportions of VISTA enhancers with enhancer-associated chromatin signatures by tissue. Active enhancers across the six considered tissues with different combinations of canonical enhancer-associated chromatin marks. For every case there are active enhancers that do not have any of these considered marks. The tissues/regions are : **(a)** forebrain, **(b)** midbrain, **(c)** hindbrain, **(d)** craniofacial, **(e)** limb, and **(f)** heart.

**Figure S4.**
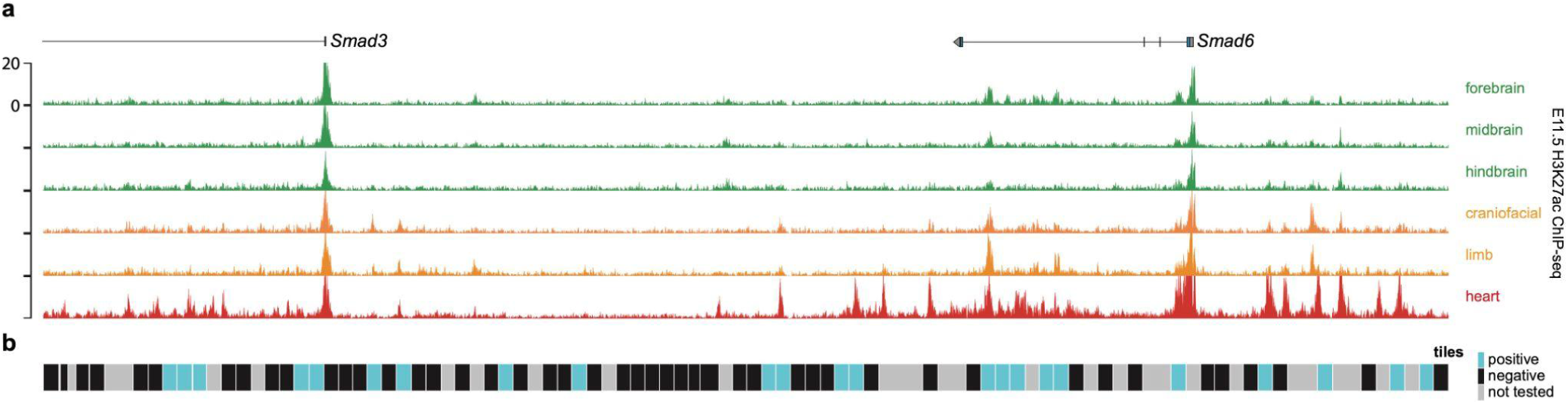
Tiling a second locus for the unbiased identification of mouse in vivo enhancers. **(a)** *Smad3*/*Smad6* locus with mouse E11.5 H3K27ac ChIP-seq data (ENCODE) for six tissues. **(b)** Elements (∼5kb in size and overlapping with adjacent elements) designed for the unbiased tiling assay. Elements that were tested and that had reproducible enhancer-reporter activity (in one or more tissues) in the mouse *in vivo* transgenic assay are shaded blue. Elements not tested are shaded gray.

**Figure S5.**
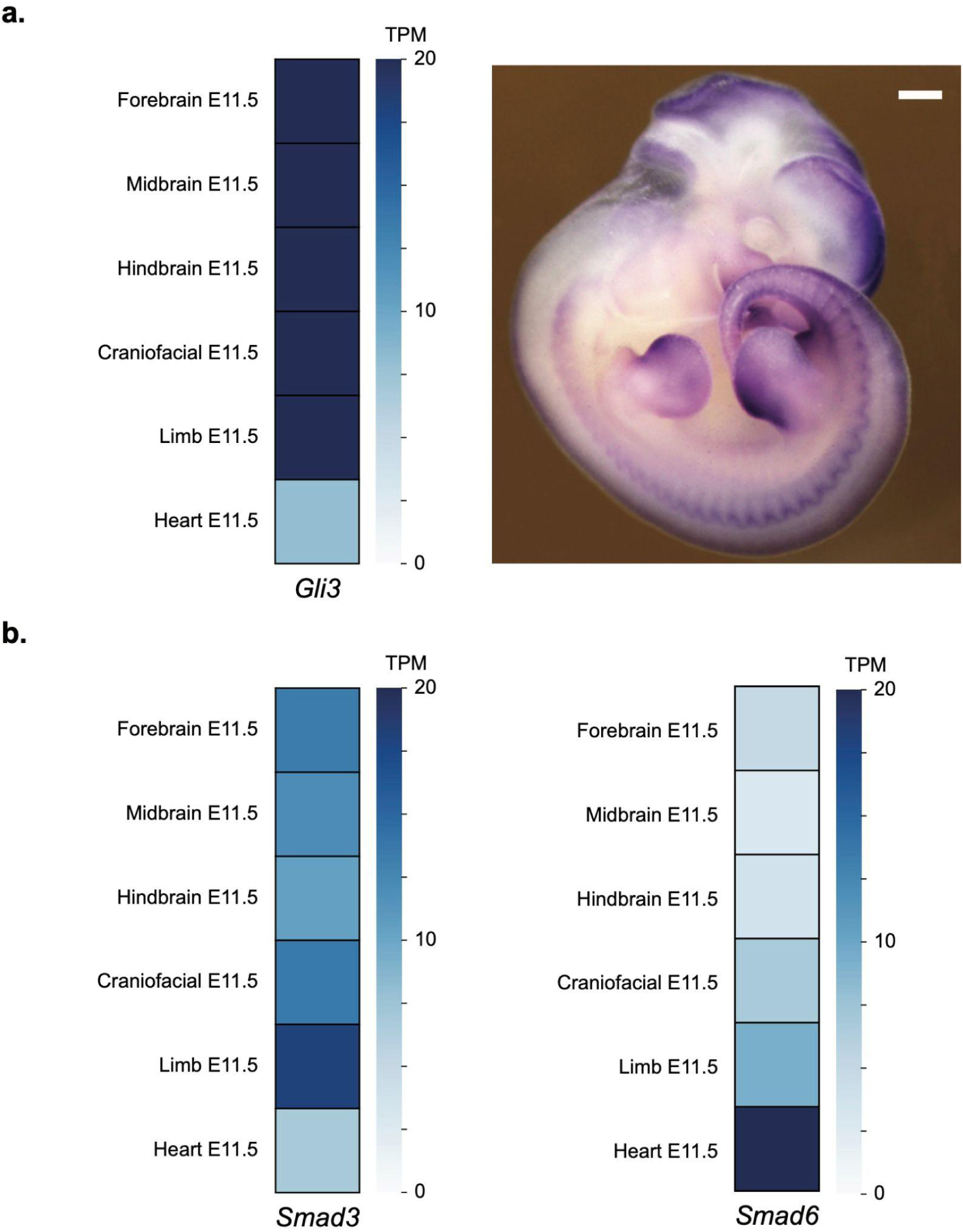
Mouse E11.5 gene expression by tissue for the *Gli3, Smad3*, and *Smad6* genes. **(a)** Per tissue RNA-seq and mouse *in situ* data for *Gli3*. **(b)** Per tissue RNA-seq data for the adjacent *Smad3* and *Smad6* genes. RNA-seq data are from mouse ENCODE^46^. TPM, transcripts per million.

**Figure S6.**
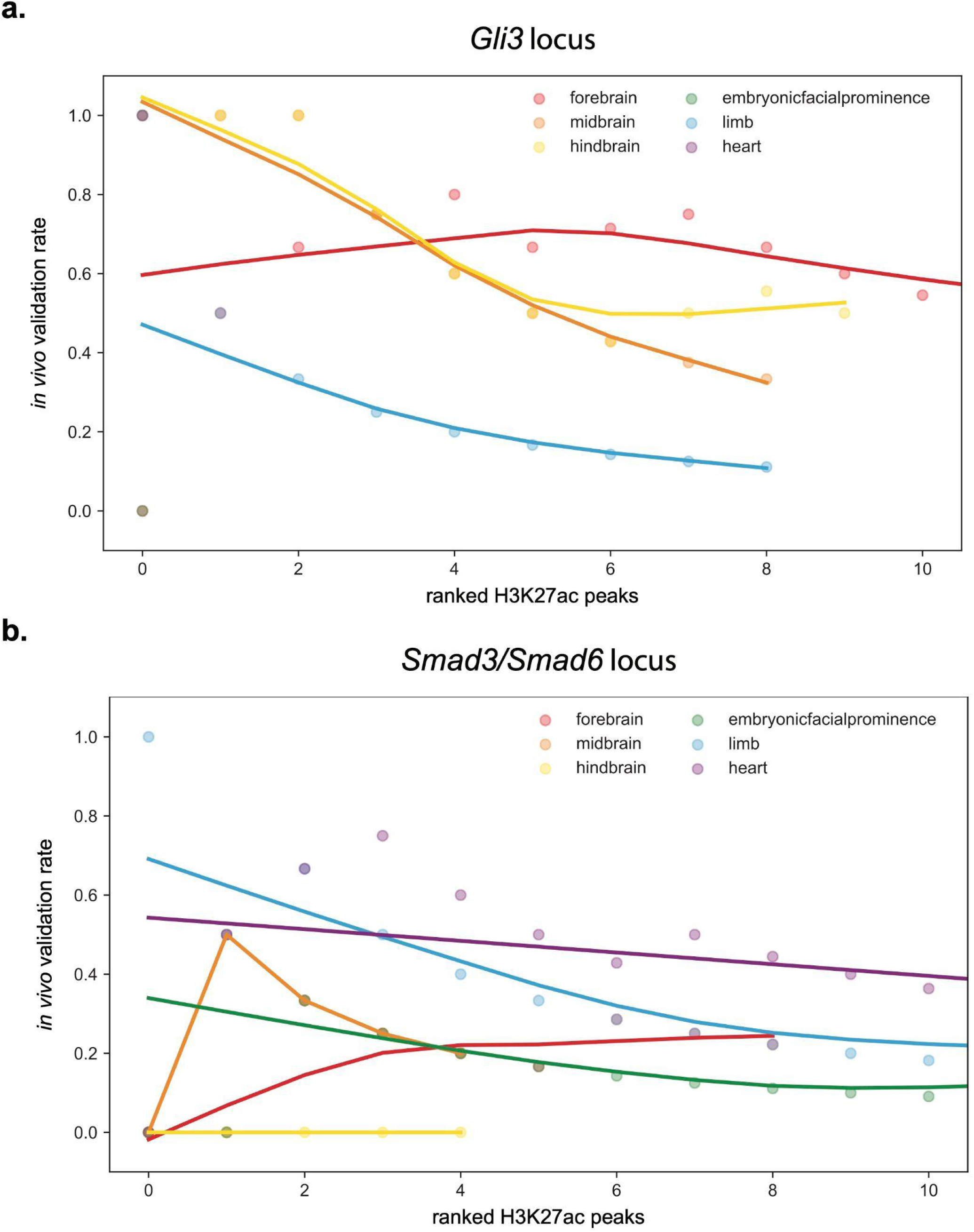
*In vivo* enhancer validation rates correlate with ranked H3K27ac peaks within tiling loci. Per tissue mouse *in vivo* enhancer validation rates of ranked H3K27ac peaks (-log_10_ q-value) for elements tested across the **(a)** *Gli3* and **(b)** *Smad3*/*Smad6* loci.

**Figure S7.**
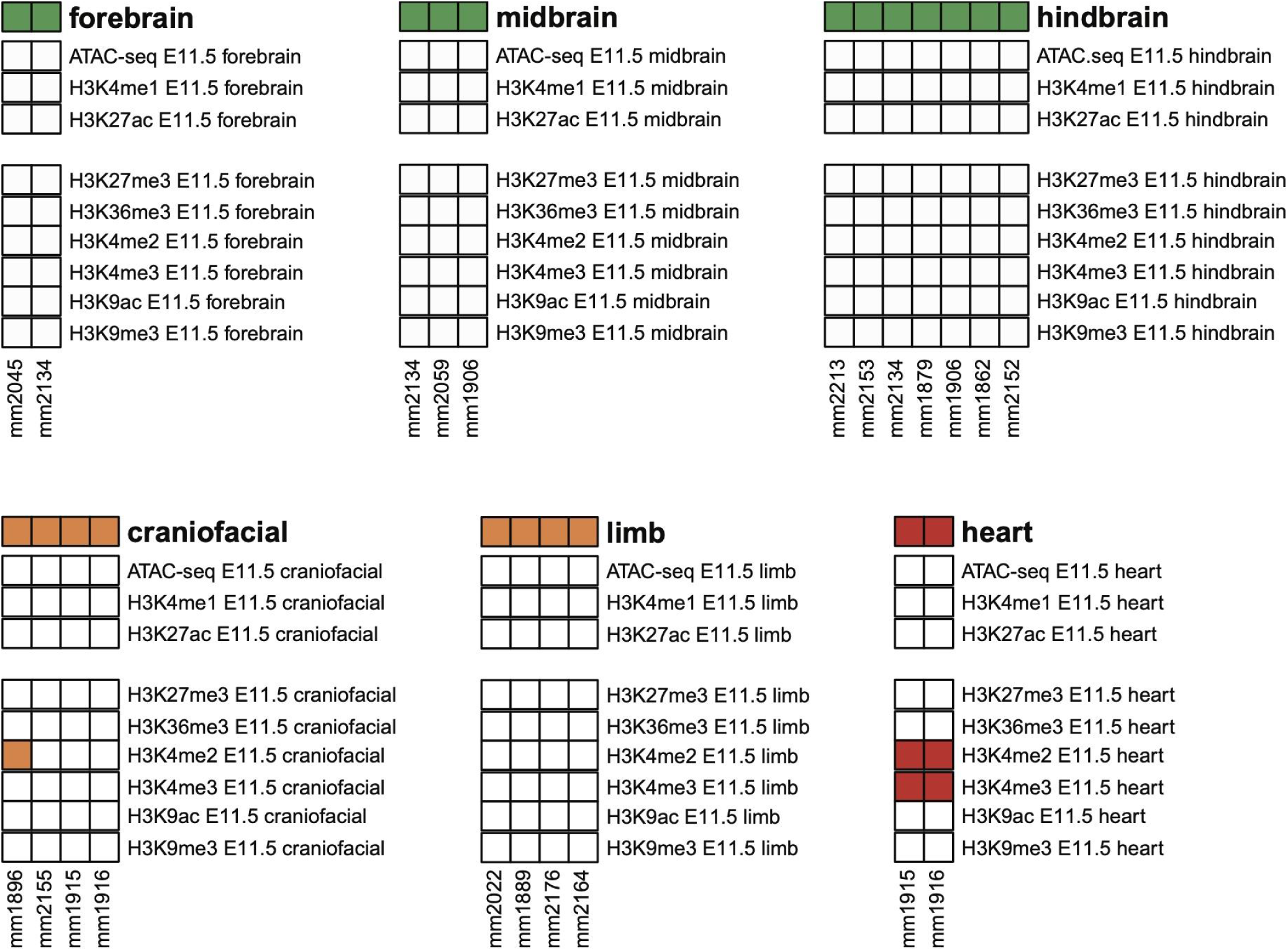
Hidden enhancers commonly lack other chromatin marks at their endogenous site. A majority of hidden enhancers identified from the unbiased tiling (across the *Gli3* and *Smad3/Smad6* loci) do not have other chromatin marks. Processed mouse chromatin data are from ENCODE^19^.

**Figure S8.**
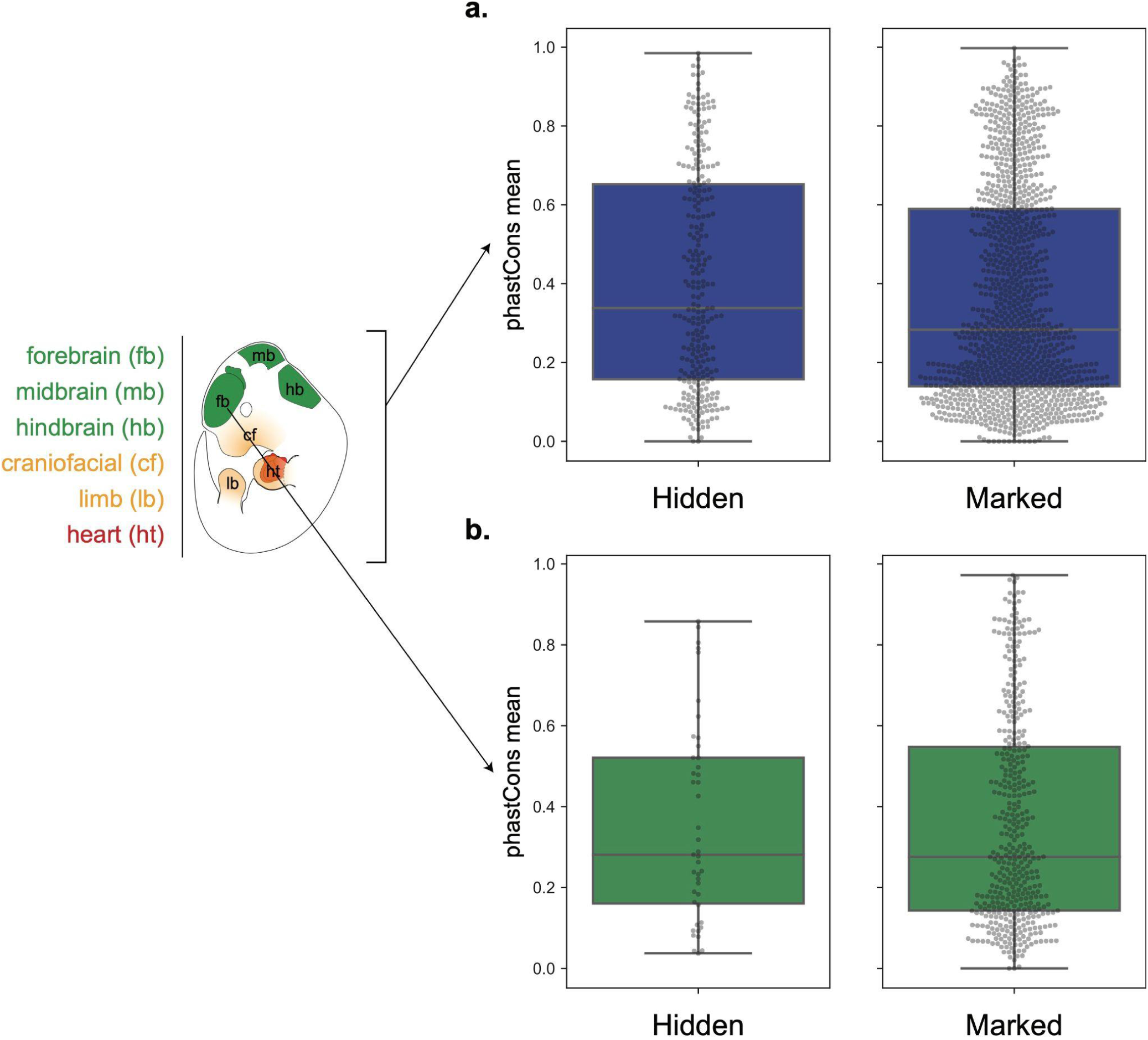
Similar levels of evolutionary conservation (phastCons) between hidden enhancers and marked enhancers. Hidden enhancers and marked enhancers have similar levels of evolutionary conservation (phastCons) for **(a)** all tissues considered together and also for each considered tissue, exemplified by **(b)** forebrain enhancers. Data not shown for the other five tissue types. No difference via Kolmogorov-Smirnov comparison.

**Figure S9.**
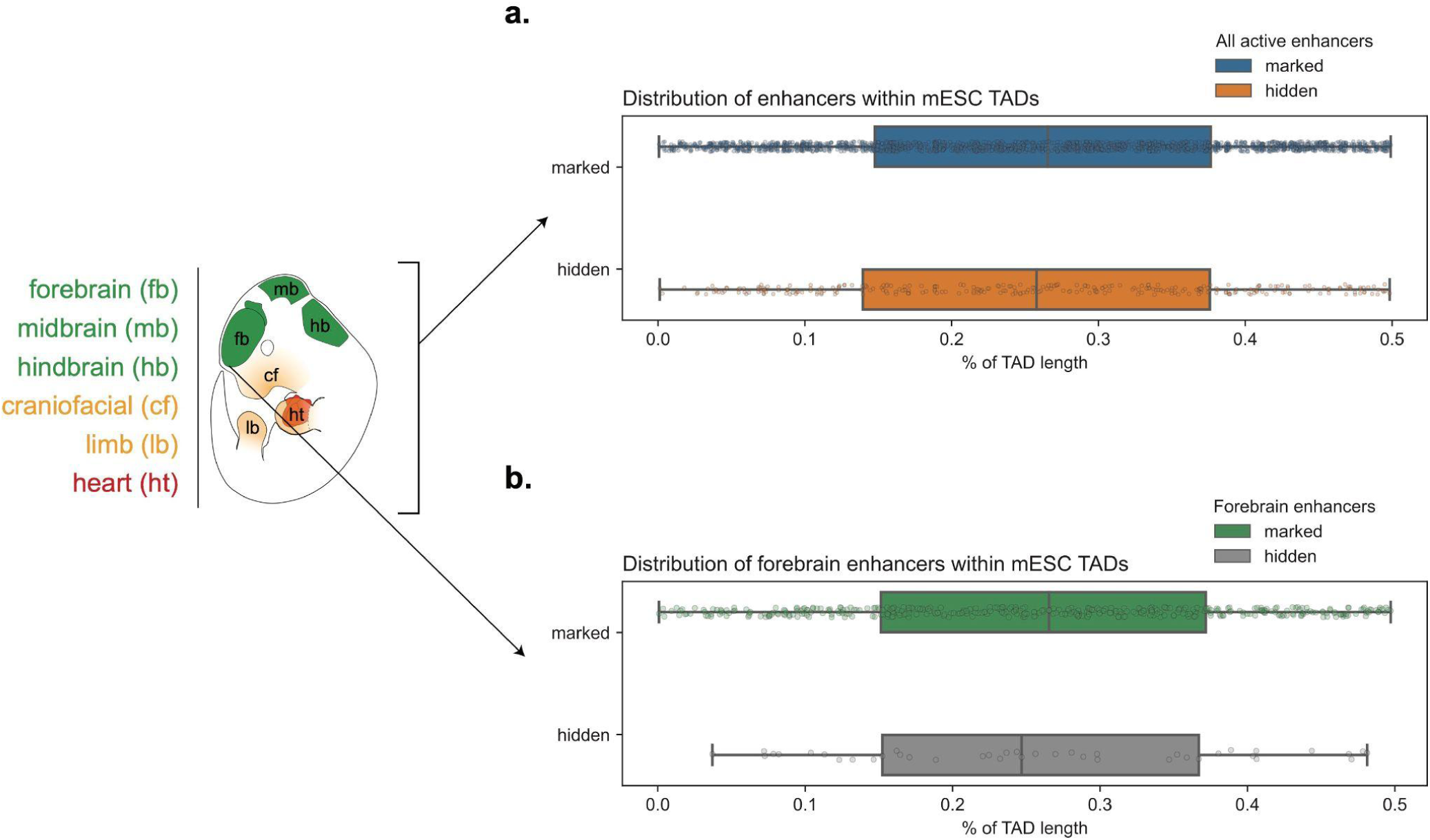
Positions of hidden enhancers within TADs are not distinguishable from those of marked enhancers. Relative to marked enhancers, hidden enhancers do not have a distinct positional bias within topologically associated domains (TADs). Summary distributions are shown **(a)** for all tissue-specific enhancers considered together and for **(b)** forebrain enhancers. Data not shown for the other five tissue types. No difference via Kolmogorov-Smirnov comparison.

## Supplementary Tables

**Table S1. ENCODE mouse chromatin and RNA-seq data**.

**Table S2. Chromatin intersections of enhancers from the VISTA retrospective study and the unbiased tiling**.

**Table S3. Overview of VISTA E11.5 enhancers by tissue**.

**Table S4. Tissue-specific H3K27ac peak counts across the two loci tested by tiling for enhancer activity**.

**Table S5. Mouse *in vivo* enhancer validation rates across the two loci tested by tiling for enhancer activity**.

**Table S6. Summary of hidden enhancer transcription factor motif analysis. Table S7. Summary of hidden enhancer functional enrichment analysis**.

## Acknowledgements

This work was supported by U.S. National Institutes of Health (NIH) grants to L.A.P. and A.V. (UM1HG009421 and R01HG003988). Research was conducted at the E.O. Lawrence Berkeley National Laboratory and performed under U.S. Department of Energy Contract DE-AC02-05CH11231, University of California (UC).

